# Type 2 Diabetes Risk Prediction Incorporating Family History Revealing a Substantial Fraction of Missing Heritability

**DOI:** 10.1101/041335

**Authors:** Jungsoo Gim, Wonji Kim, Soo Heon Kwak, Kyong Soo Park, Sungho Won

**Affiliations:** Institute of Health and Environment, Seoul National University; Interdisciplinary Program of Bioinformatics, Seoul National University; Department of Internal Medicine, Seoul National University College of Medicine; Graduate School of Public Health, Seoul National University

## Abstract

Despite many successes of genome-wide association (GWA) studies, known susceptibility variants identified by GWAS have the modest effect sizes and we met noticeable skepticism about the risk prediction model building with large-scale genetic data. However, in contrast with genetic variants, family history of diseases has been largely accepted as an important risk factor in clinical diagnosis and risk prediction though; complicated structures of family history of diseases have limited their application to clinical use. Here, we develop a new method which enables the incorporation of general family history of diseases with the liability threshold model and a new analysis strategy for risk prediction with penalized regression incorporating large-scale genetic variants and clinical risk factors. An application of our model to type 2 diabetes (T2D) patients in Korean population (1846 cases out of 3692 subjects) demonstrates that SNPs accounts for 28.6% of T2D’s variability and incorporation of family history leads to additional improvement of 5.9%. Our result illustrates that family history of diseases can have an invaluable information for disease prediction and may bridge the gap originated from missing heritability.

## INTRODUCTION

Even though some significant results from genome-wide association studies (GWAS) have been successfully translated into clinical utility^1^, many studies showed that genetic screening for the prediction of complex diseases had currently little value in clinical practice^2^. For example, heritability estimates of type 2 diabetes (T2D) from twin and familial studies ranged from 40% to 80%^3,4^. However, the estimated proportions of heritability explained by known susceptibility variants of T2D have been from 10% to 27.93%, and it indicates that most heritability is still unexplained^5^^-^^7^. In addition to this so-called ‘missing-heritability’ issue, GWAS-based common variants tend to mildly predispose to common disease^8^, which generates some doubt about clinical utility of GWAS findings to risk assessment in clinical care^9^.

Alternatively, family history reflects genetic susceptibility, and also interactions between genetic, environmental, cultural, and behavioral factors^10,11^. Therefore, it has been repeatedly addressed that the incorporation of family history of diseases to the risk prediction model might implicitly cover effects of uncovered genetic risk factors and shared gene-environment interaction^12,13^, and thus it has been often expected as an important risk factor in clinical assessment^13^.

There have been many investigations for disease risk prediction with large-scale genetic data and family history of diseases. Most popular approaches for disease risk prediction are based on logistic regression with genotype scores. With train set, regression coefficients of some significantly associated SNPs^14^ are calculated and sums of the weighted genotype scores with their regression coefficients are incorporated as a single covariate to the logistic regression for test set^15^. However the accuracy of such disease risk prediction models has been much lower than that of expected from the heritability estimates. To overcome the controversy over potential clinical usage of GWAS findings, several approaches have been proposed to include a large number of SNPs into the prediction model: using penalized regression methods^16,17^ and random effects model^18^. However, these attempts still have several limitations. For penalized approaches, computational intensity linearly or quadratically increases with the number of SNPs^16^ and thus the accuracy of the prediction model with penalized regression depends on the initial feature screening step because certain number of SNPs has been chosen from the marginal effects of SNPs and joint effects of SNPs are ignored for feature selection. Speed et al solved this problem with a random effect model for linear regression where disease statuses are considered as continuous response variable. In such a case the substantial bias can be observed if the probability of being affected is very small or large^18^.

In this report, we propose a new disease risk prediction model with penalized regression with following features: (i) a certain number of SNPs is selected with best linear unbiased prediction, (ii) conduct the penalized logistic regression analyses using both SNPs and clinical variables, and (iii) provide a new method to incorporate the general family history of diseases. However, in spite of their importance, familial relationships of relatives with known disease statuses are usually heterogeneous between subjects, and thus they were limitedly utilized for disease prediction model. An application of our model to type 2 diabetes (T2D) patients in Korean population (1846 cases out of 3692 subjects) demonstrates that SNPs accounts for 28.6% of T2D’s variability and incorporation of family history leads to additional improvement of 5.9%. Our result illustrates that family history of diseases can have an invaluable information for disease prediction and may bridge the gap originated from missing heritability.

## METHODS

### Evaluating posterior mean of disease risk of an subject using family history

We assume that genotypes are not used to estimate posterior mean of disease risk and environmental effects are known. We started our model by evaluating posterior mean of disease risk using the standard liability threshold model^19^. We assume that disease statuses are determined by the unobserved liabilities (denoted as *L*) and if they are larger than a threshold *T*, which is determined by the prevalence, he/she becomes affected and they are normally distributed. In this section, we let ***Y***_*i*_ = (*Y*_*i*_0__,*Y*_*i*_1__,…,*Y*_*i*_*n*-1__)^*t*^, ***L***_*i*_ = (*L*_*i*_0__,*L*_*i*_1__,…,*L*_*i*_*n*-1__)^*t*^, and ***Z***_*i*_ = (*Z*_*i*_0__,*Z*_*i*_1__,…,*Z*_*i*_*n*-1__)^*t*^ respectively represents phenotypes, liabilities, and environment vectors of the subject *i* and his/her family in order. We used subscript *i*_*j*_ to indicate each family member of the subject *i* (*j* = 0 indicates the subject *i* itself). We further denote by *ƒ*_*j*_ and *ψ*_*jj*′_ the inbreeding coefficient for relative *j* of the subject *i* and the kinship coefficient between two relatives *j* and *j*′ of the subject *i*, respectively. It should be noted that *ψ*_*jj*′_ is 0 if the subjects *j* and *j*′ are in different families. We then define the kinship coefficient matrix as **Ψ**_i_ where (**Ψ**_i_)_*jj*′_ is 2*ψ*_*jj*′_ for *j* ≠ *j*′, and 1 + *ƒ*_*j*_ otherwise. With this notation, we assumed that

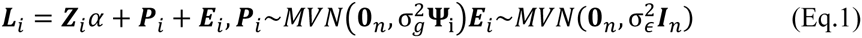

where ***I***_*n*_ is *n* × *n* dimensional identity matrix, **0**_*n*_ and **1**_*n*_ are *n* dimensional column vectors. Here 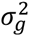 and 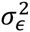 indicate the variances of polygenic effect and random effect, respectively.

Based on this liability threshold model, we can calculate the conditional expectation of *L*_*i*_, PM, when the family histories of disease are conditioned. We let the subscript *i*_*j*_ indicates relative *j* of the subject *i*. We further define a random variable ***A***_*i*_ of the subject *i* by

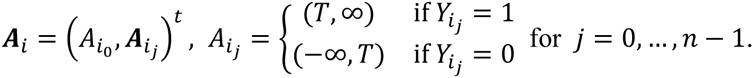

let *I*_*A*_*i*_*j*___ (*L*_*i*_*j*__) = 1 if *L*_*i*_*j*__ ∈ *A*_*i*_*j*__ and otherwise 0, and I_*A*_*i*__(***L*_*i*_**) = (I_*A*_0__(*L*_*i*_0__), …,I_*A*_*n*-1__(*L*_*n*-1_))^*t*^, then PM becomes

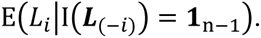

PM can be calculated with the moment generating function (mgf) of truncated multivariate normal distribution to calculate the conditional distribution. The joint probability density function (pdf) can be defined as

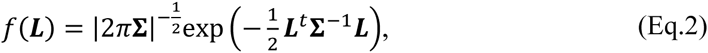

where **Σ** = cov(***L***). Based on the conditional pdf of ***L*** given I(***L***) = **1** and some algebra, we can have

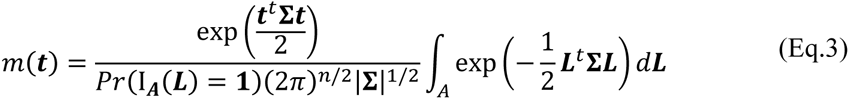

If we let (Σ)_*jk*_ = σ_*jk*_ and *F*_*k*_(x) be the marginal pdf of *L*_*k*_, the PM for subject *i* can be obtained by

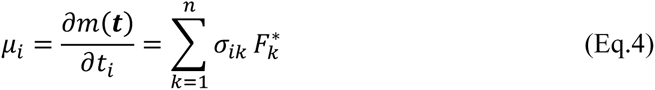

where

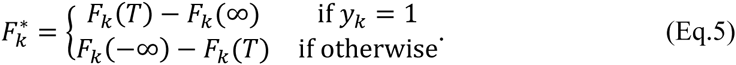

Derivation of *F*_*k*_ requires marginal pdf of truncated multivariate normal distribution, and it can be derived as follow. First, we partitioned ***L*** into two parts *L*_*i*_ and ***L***(-*i*) and then ***L*** can be rewritten as,

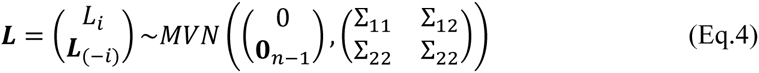

If we denote the lower and upper truncated point of ***L*** as ***a*** and ***b*** respectively, then the truncated normal distribution function when ***a*** < *L* < ***b*** becomes

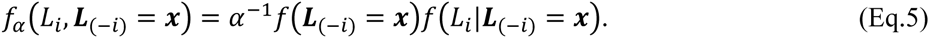

By using the marginal pdf of ***L***_(-*i*)_ at ***L***_(-*i*)_ = ***x*** and the fact that conditional distribution of normal distribution is normally distributed, one can easily show that *L*_*i*_|***L***_(-*i*)_ = ***x*** follows normal distribution with *E*(*L*_*i*_|***L***_(-*i*)_ = *x*) = 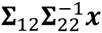 and *Var*(*L*_*i*_|***L***_(-*i*)_ = ***x*** = 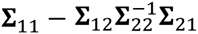. With these results, the multivariate marginal pdf of ***L***_(-*i*)_ becomes

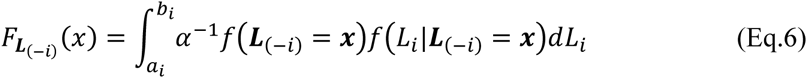

The integral can be readily computed by using conventional statistical software and we used *pmvnorm()* function in R package *mvtnorm*^20^.

### Prescreening with best linear unbiased predictor

To select an effective list of SNPs, we considered the best linear unbiased prediction (BLUP) of SNP effects using GCTA^21^. GCTA provides the BLUP of total genetic effect for all subjects by considering a mixed linear model with random effects of SNPs, i.e., ***y*** = ***x***_***Z***_***β*** + ***g*** + ***ϵ*** with *var*(***y***) = ***V*** = 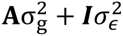, where ***y*** and ***β*** are a vector of phenotypes and fixed effect of subjects with genotypes, respectively, and ***g*** and ***ϵ*** are vectors of total genetic effects of the subjects with 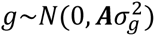 and and residual effects with 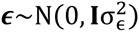. **A** is the genetic relationship matrix (GRM) between subjects. By estimating GRM from all the SNPs, the BLUP of ***g*** can be provided by the restricted maximum likelihood (REML) approach.

Consider a mathematically equivalent model, ***y*** = ***x***_***Z***_***β*** + ***Wu*** + ***ϵ*** with *var*(***y***) = ***V*** = 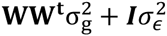, s, where ***u*** is a vector of random effects with 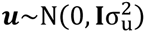 and ***W*** is a standardized genotype matrix. The GRM, **A**, can be defined by ***WW***^***t***^/*p*_1_, where *p*_1_ is the number of SNPs. Since these two equations are mathematically equivalent, the BLUP of ***g*** can be transformed to the BLUP of ***u*** by 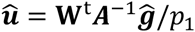. Thus the estimate of *u*_*i*_ corresponds to the coefficient *w*_*iG*_*l*__ which is the *G*_*l*_th SNP of the *i*th subject element of **W**. Note that 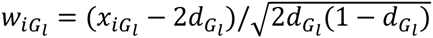, where *x*_*iG*_*l*__ and *d*_*G*_*l*__ are the numbers of copies of the reference allele and the frequency of the reference allele, respectively. Divided by 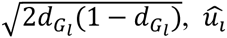 can be rescaled for the original genotype.

### Penalized regression method

We let ***x***_***i***_ = (***x***_*iG*_, ***x***_*iZ*_) and *y*_*i*_ be a covariate vector and a dichotomous phenotype for the subject *i*, and affected and unaffected subjects are coded as 1 and 0, respectively. We further denote *x*_*iG*_*l*__ and *x*_*iZ*_*m*__ be a coded genotypes of the *l*th SNP and the *m*th clinical covariate, respectively. The *p* dimensional coefficient vector ***β*** = (*β*_1_,…,*β*_*p*_)^*t*^ consists of *p*_1_ genetic variants and *p*_2_ clinical variables. Under this model, ***β*** can be estimated by minimizing the penalized negative log-likelihood:

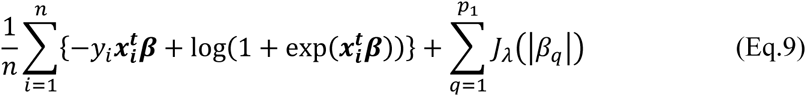

where *J*_*λ*_ is a penalty function and *λ* is a vector of tuning parameter that can be determined by a search on an appropriate grid. Note that only genetic variants were penalized in Eq. 9.

With the different choice of penalty function, lasso^22^, ridge^23^, EN^24^, SCAD^25^ and TR^26^ can be performed. The penalty of Lasso is *J*_*λ*_(*t*) = *λt* and it has been often utilized because Lasso can conduct both shrinkage and variable selection. Even though lasso has an overfit problem, it shows a quite stable performance especially then sample size is small. Ridge uses *J*_*λ*_(*t*) = *λt*^2^ as its penalty. Similar to lasso, it has shrinkage effect by choosing *λ* but no selection of variables. Ridge can be conducted even when *p* is much larger than *n*. EN, which is a convex combination of lasso and ridge, has a penalty of *J*_*λ*_(*t*) = *λ*(*αt* + (1 - *α*)*t*^2^), and we considered 20 equally spaced grid points from zero to one for *α*. EN enables us to have balanced estimates, producing a slightly more complex model than lasso but far simpler model then ridge. The penalty of SCAD is 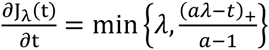 and we used a = 50 for our own optimization algorithm. SCAD is known to have the oracle property, i.e., the set of selected variables are asymptotically equal to the set of true causal variables. In spite of the theoretical optimality, SCAD estimates can be poor unless the sample size is large and the effects of signal variables are strong. For TR estimates, we first obtained ridge estimates with tuning parameter *λ* and then truncated them with a level *a*, making coefficients whose absolute values smaller than *a* as zero. For the appropriate choice of truncating level, 20 grid points equally spaced in logarithmic scale from minimum to maximum ridge estimates were considered for *a*. All the analysis was performed with *glmnet*^27^ R package.

### Building disease risk model using penalized regression method

In this section, we describe how we developed a disease risk model with the estimated PM score. Followings are the brief steps.

1. We consider Age, Sex, BMI, SBP, and DBP as clinical covariates, and they are included for all regression.
2. Calculate PM for all subjects with family histories of diseases.
3. We conduct 10 fold cross validation. That is, we divide dataset into 10 different sub-data, and one and the other nine subdata are used as test and train set respectively.
4. Using train set, we select *k* SNPs with pvalues about the marginal effects of SNPs from logistic regression, and the proposed BLUP method. We considered k = 100, 500, 1000, 5000, 10000, 20000.
5. Perform Lasso^22^, Ridge^23^, Elastic-Net24 (EN), SCAD^25^ and Truncated Ridge^26^ (TR) for penalized regression and mixed effect model (MultiBLUP^18^). Tuning parameters for each penalized regression are chosen with additional 10 fold cross-validation with train set. We divide train set into 10 different subdata, and for different choices of tunning parameter, we get the prediction model with 9 subdata. Then calculate the AUC with the remaining 1 subdata, and tunning parameters which result in the largest AUC are finally chosen.
6. The prediction models for penalized regressions and multiBLUP are applied to the test set, and we calculate AUCs.
7. Repeat 3-7 for the different combinations of train and test set

### Data Description

To demonstrate the validity of our proposed model and to illustrate its application to risk prediction, we investigated two real datasets: KARE and SNUH. Since SNUH dataset has cases only, we merged two datasets by adjusting platform difference (matching SNPs existing in both platforms and imputing NAs using Shapeit). Briefly, we analyzed 3692 subjects (1846 cases / 1846 controls) with 267,063 SNPs.

KARE cohort was collected to construct an indicator of disease of genetic character in an attempt to predict outbreaks of diseases. There are initially 8,842 participants and they were genotyped for 352,228 SNPs with the Affymetrix Genome-Wide Human SNP array 6.0. In our study, the following SNPs were discarded in further analysis: (1) p-values for Hardy-Weinberg equilibrium (HWE) are less than 10^−5^, (2) genotype call rates are less than 95% and (3) MAFs are less than 0.05. We also eliminated subjects with gender inconsistencies, whose identity in state (IBS) were more than 0.8 or whose call rates were less than 95%. Participants were asked whether they have affected relatives and if so, their ages and familial relatedness. These family histories of diseases including T2D are also available for KARE data. Finally, 1,167 T2D cases and randomly selected 1846 controls with 267,063 SNPs were used for the analysis.

For SNUH data, T2D patients were diagnosed as T2D using the World Health Organization criteria for Seoul National University Hospital, and 681 subjects with positive family history of diabetes in the first-degree relatives were preferentially included. The family history of their relatives was based on the recall of the proband. However, family members were encouraged to perform a 75 g oral glucose tolerance test, and subjects positive for glutamic acid decarboxylase autoantibodies test were excluded. In total, the disease statuses of 7,825 relatives were available and among them 2,875 subjects had T2D. T2D patients originally diagnosed from Seoul National University Hospital were genotyped with the Affymetrix Genome-Side Human SNP array 5.0, and 480,589 SNPs were obtained. The same conditions for quality control with KARE were applied, two subjects and a number of SNPs were excluded. In total, 679 T2D patients with 267,063 SNPs were used for the analysis.

### Estimating variability in penalized logistic regression

To estimate the variability of each variable in the penalized regression model, we used residual deviance from the penalized log-likelihood. The residual deviance is defined as,

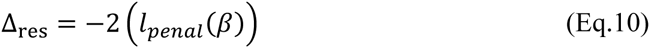

where 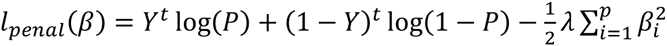 and 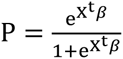 Using eq.10, we defined variability explained by *i*th reduced model as

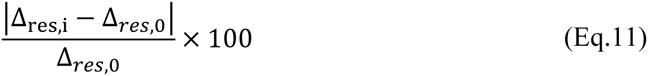

where Δ_*res*,0_ denotes the residual deviance of the null model.

## RESULTS

### Characteristics of the variables

As described previously, established a methodology for estimating the PM for all subjects in a pedigree and applied the method to real dataset. As can be seen in the Fig. 1A, mean values of PM between T2D cases and controls were not distinct. However, more subjects with T2D have high PM (larger than 0.5) compared to control subjects. Boxplot of other clinical covariates between cases and control are shown (Fig. 1).

**Figure 1.**
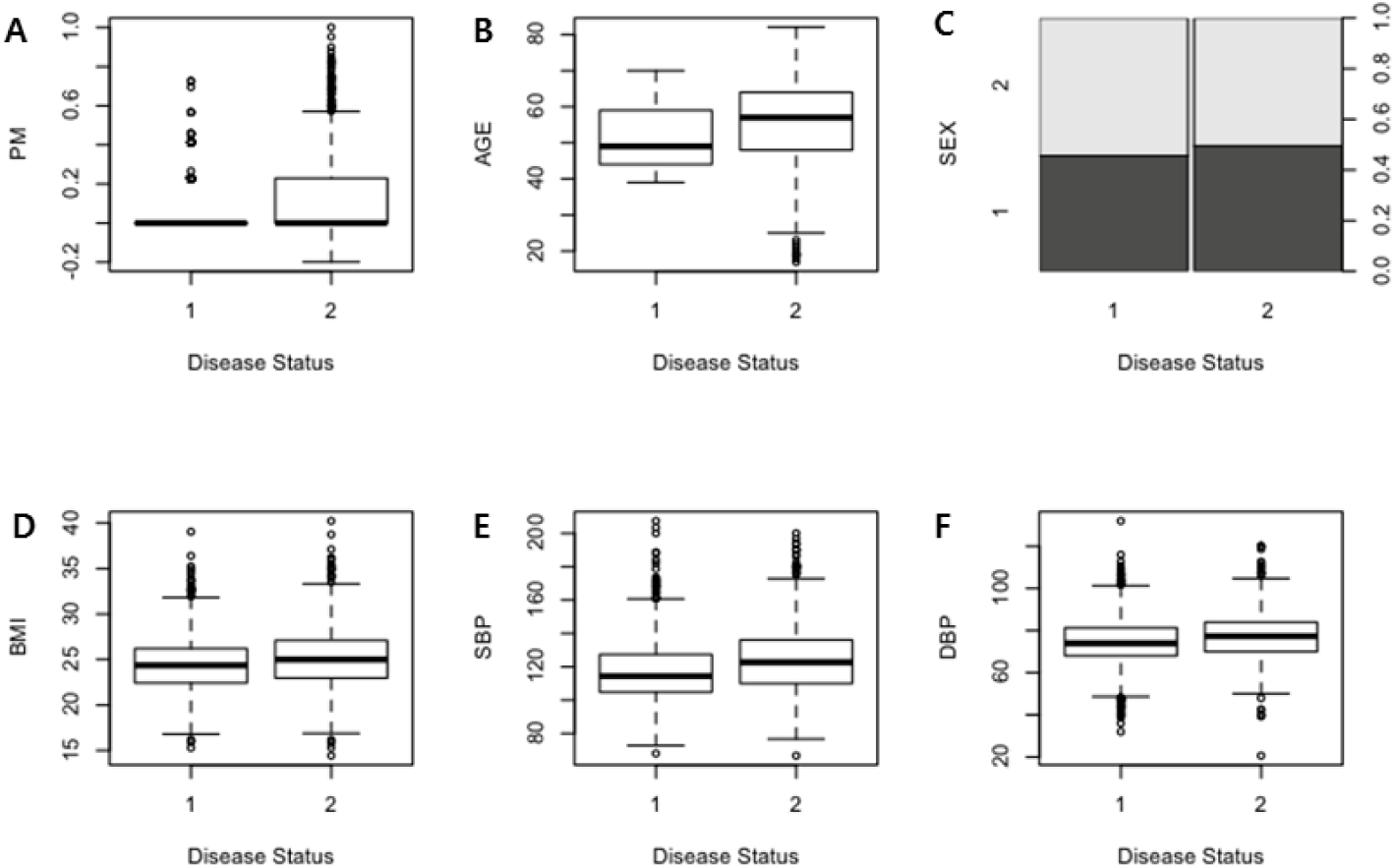
Characteristics of variables. Characteristics of the PM (A), age (B), sex (C), BMI (D), SBP (E) and DBP (F) are shown in boxplots. Here disease status 1 and 0 indicates T2D case and control, respectively.

To find the most effective set of SNPs, we selected SNPs based on p-value obtained from logistic regression and BLUP obtained by mixed effect model. Since the selected set of SNPs should be applied in penalized regression, we expected it would be more effective if the set of SNPs uniformly distributed across the genome. We discretized whole genome with a window size of 5M base pair and counted the frequency of SNPs in each window. With varying number of SNPs (0.1k to 20k), it is apparent that both set of SNPs selected by p-value and BLUP criteria exhibit similar patterns (Fig. 1G).

### A comparison of performances

The main purpose of this work was to construct a T2D risk prediction model. To find the best model, we sought to compare the performances of six methods with different criteria of selecting SNPs and varying number of SNPs. We repeated our analysis with family history Table I) and without family history (Table II). On the whole and the most interestingly, family history (PM) plays a very important role in risk prediction for all methods except MultiBLUP. By comparing Table I and II, it is obvious that a significant improvement was obtained with a prediction model using PM variables. A striking example can be seen in truncated ridge with 5000 SNPs selected by BLUP criteria (Table I and II), changing AUC from 0.689 to 0.736.

**Table I.** AUC with clinical variables and SNPs

| CRITERIA | # of SNPs | RIDGE | LASSO | EN | SCAD | T.RIDGE | MultiBLUP |
| --- | --- | --- | --- | --- | --- | --- | --- |
| P-value | 100 | 0.642 | 0.637 | 0.637 | 0.616 | 0.641 | 0.532 |
|  | 500 | 0.640 | 0.626 | 0.626 | 0.608 | 0.640 | 0.542 |
|  | 1,000 | 0.640 | 0.624 | 0.624 | 0.608 | 0.640 | 0.544 |
|  | 5,000 | 0.660 | 0.635 | 0.635 | - | 0.660 | 0.546 |
|  | 10,000 | 0.668 | 0.640 | 0.640 | - | 0.668 | 0.560 |
|  | 20,000 | 0.674 | 0.640 | 0.640 | - | 0.674 | 0.582 |
| BLUP | 100 | 0.611 | 0.602 | 0.602 | 0.585 | 0.612 | 0.500 |
|  | 500 | 0.614 | 0.600 | 0.600 | 0.594 | 0.614 | 0.513 |
|  | 1,000 | 0.626 | 0.611 | 0.611 | 0.601 | 0.626 | 0.537 |
|  | 5,000 | **0.689** | 0.647 | 0.647 | - | **0.689** | 0.581 |
|  | 10,000 | 0.672 | 0.626 | 0.626 | - | 0.672 | 0.550 |
|  | 20,000 | 0.674 | 0.639 | 0.639 | - | 0.674 | 0.571 |

**Table II.** AUC with clinical variables, SNPs and PM

| CRITERA | # of SNPs | RIDGE | LASSO | EN | SCAD | T.RIDGE | MultiBLUP |
| --- | --- | --- | --- | --- | --- | --- | --- |
| P-value | 100 | 0.693 | 0.687 | 0.688 | 0.676 | 0.693 | 0.534 |
|  | 500 | 0.687 | 0.672 | 0.672 | 0.665 | 0.687 | 0.544 |
|  | 1,000 | 0.685 | 0.669 | 0.669 | 0.664 | 0.685 | 0.536 |
|  | 5,000 | 0.709 | 0.687 | 0.687 | - | 0.709 | 0.541 |
|  | 10,000 | 0.717 | 0.690 | 0.690 | - | 0.717 | 0.554 |
|  | 20,000 | 0.721 | 0.689 | 0.689 | - | 0.721 | 0.561 |
| BLUP | 100 | 0.669 | 0.659 | 0.659 | 0.643 | 0.669 | 0.500 |
|  | 500 | 0.659 | 0.642 | 0.642 | 0.639 | 0.659 | 0.505 |
|  | 1,000 | 0.670 | 0.651 | 0.651 | 0.645 | 0.670 | 0.516 |
|  | 5,000 | **0.736** | 0.691 | 0.691 | - | **0.736** | 0.575 |
|  | 10,000 | 0.721 | 0.673 | 0.673 | - | 0.721 | 0.544 |
|  | 20,000 | 0.725 | 0.689 | 0.689 | - | 0.725 | 0.562 |

In the majority of cases, truncated ridge and ridge revealed a higher prediction performance. Interestingly, similar behavior was observed between ridge and truncated ridge, and between lasso and elastic net. For a small number of SNPs, p-value criteria showed better performance. However, the difference gets negligible (even reversed) as the number of SNPs increased.

The best performance (AUC = 0.736) was observed in ridge and truncated ridge with PM and 5000 SNPs selected by BLUP criteria (truncated ridge showed slightly higher AUC O(E-05). This is consistent with results obtained in previous studies.^28,29^ To investigate the effect of each variables, we built the logistic regression without any SNPs. Based on the nested 10 fold cross validation scheme, which was applied in our model building steps, we measured the performance of logistic model without PM and with PM. Without PM, the AUC value was 0.672, but increased to 0.730 with PM included (Table III). This value is not much different from the highest AUC (0.736) obtained with 5000 additional SNPs.

**Table III.** AUC without SNPs

| VARIABLES INCLUDED | LOGISTIC REGRESSION |
| --- | --- |
| AGE, SEX, SBP, DBP, BMI | 0.672 |
| AGE, SEX, SBP, DBP, BMI, PM | 0.730 |

We measured the time complexity of each method. Table IV shows the result. In general, the analysis time increased if the number of SNPs increased, except MultiBLUP. In case of MultiBLUP, it has several manual steps to perform a prediction analysis. Therefore, it was difficult to measure exact time for the whole analysis steps. However, MultiBLUP was not affected much by the number of SNP increment.

**Table IV.** Analysis Time

| # of SNPs | RIDGE | LASSO | EN | SCAD | T.RIDGE | MultiBLUP |
| --- | --- | --- | --- | --- | --- | --- |
| 100 | 15.6 sec | 13.2 sec | 4.7 min | 37 min | 1.9 min | < 20 min |
| 500 | 1.2 min | 1.2 min | 25.1 min | 5.2 hour | 6.0 min | < 20 min |
| 1,000 | 2.6 min | 2.2 min | 43.5 min | 12.2 hour | 11.1 min | < 20 min |
| 5,000 | 12.3 min | 53.7 min | 35.6 min | ~ 3 days* | 34.4 min | < 20 min |
| 10,000 | 24.3 min | 1.7 hour | 1.1 hour | ~ 6 days* | 1.7 hour | < 20 min |
| 20,000 | 47.7 min | 3.4 hour | 3.4 hour | ~ 12 days* | 3.3 hour | < 20 min |

\*Not measured but estimated

### Variability explained by each variable

To estimate the variability explained by each variable, we investigated the model with 5000 SNPs selected by BLUP. As described in the method section, we fitted the several reduced model to evaluate the residual deviance of each variable. Figure 2 illustrates the findings of this analysis. The largest portion (58.9%) remained unexplained, indicating the variables in the model is not good enough to explain the data. The second largest portion (28.6%) was from the SNPs. Even though the prediction performance was not significantly increased with these SNPs, they explained about 30% of variability. In contrast, PM which showed dramatic increase in prediction AUC, explained only 5.9% of total variability.

**Figure 2.**
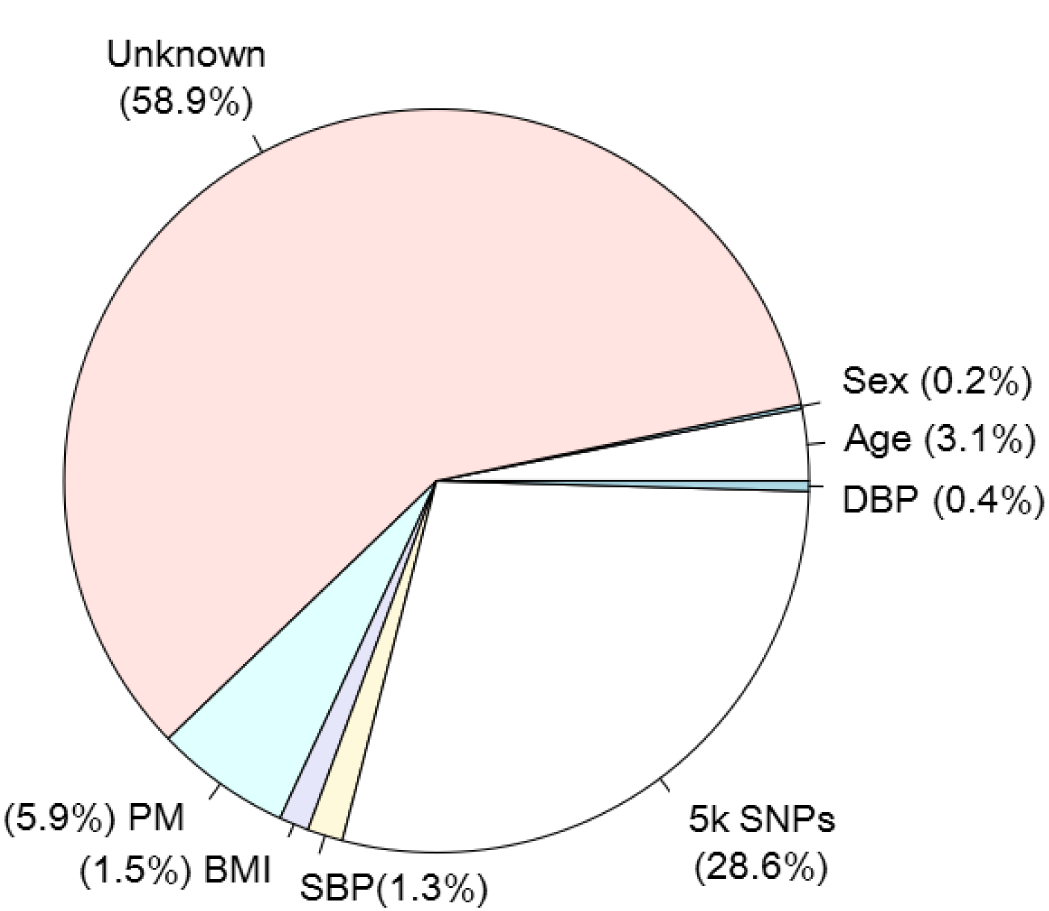
Variability pie chart. Variability explained by each variable in the final model is shown. For six clinical variables (Age, Sex, BMI, SBP, DBP, PM), variability is shown with its own proportion, while the variabilities of 5,000 SNPs is shown with their summed proportion.

## DISCUSSION & CONCLUSIONS

Prior works have documented the effectiveness of combining many SNPs using regularization methods or incorporating family history in improving prediction performance of disease risk^10,11,16^. However, these studies have either been one-sided studies or not simultaneously focused on both sides: combining more SNPs and incorporating family history. In this study we tested the extent to which combining SNPs and incorporating family history improved risk prediction with a group of T2D patients and controls. For that purpose we first developed a method estimating the posterior mean of being affected for the subjects in a pedigree. Then we compared the prediction performance of six different methods using SNPs selected by p-value obtained from logistic regression and BLUP obtained from mixed effect model. To more reliably validate the model, we performed the nested cross-validation scheme. Even though it is time-consuming, known to be more reliable.

What we found in this study is that in virtually all cases, including family history (evaluated as PM) greatly improved the prediction performance while SNPs showed slight improvement. These findings extend those without SNPs, confirming that family history tends to produce more effective genetic or environmental effect on prediction result than on SNPs. This study, therefore, indicates that the benefits gained from including PM may address a need for finding gene-gene interaction or gene-environmental interaction effects across a wide range of complex diseases.

However, some limitations worth noting. One of the limitations of our study is that we did not consider other types of structural variants, such as copy number variation, which might affect a risk of T2D but the contribution is poorly known. It is more recommendable to include rarer risk alleles with large effects and gene-gene, or gene-environment interaction into the prediction model. More of the genetic risk can be explained as more causal risk variants are identified. However, rare variant analyses or interaction analyses require more complicated statistical methods to effectively analyze the effects. Therefore the ultimate goal of the future work is to integrate advanced statistical methods with genetic data and biological knowledge, which will further improve the power to detect complex interactions efficiently.

## ACKNOWLEDGEMENTS

This research was supported by Basic Science Research Program through the National Research Foundation of Korea (NRF) funded by the Ministry of Education (NRF-2013R1A1A2010437) and also supported by the National Research Foundation of Korea Grant funded by the Korean Government (NRF-2014S1A2A2028559).

